# Recognition of centromere-specific histone Cse4 by the inner kinetochore Okp1-Ame1 complex

**DOI:** 10.1101/2023.06.06.543957

**Authors:** Sunbin Deng, Jiaxi Cai, Stephen C. Harrison, Huilin Zhou, Stephen M. Hinshaw

## Abstract

Successful mitosis depends on the timely establishment of correct chromosomal attachments to microtubules. The kinetochore, a modular multiprotein complex, mediates this connection by recognizing specialized chromatin containing a histone H3 variant called Cse4 in budding yeast and CENP-A in vertebrates. Structural features of the kinetochore that enable discrimination between Cse4/CENP-A and H3 have been identified in several species. How and when these contribute to centromere recognition and how they relate to the overall structure of the inner kinetochore are unsettled questions. We have determined the crystal structure of a Cse4 peptide bound to the essential inner kinetochore Okp1-Ame1 heterodimer from budding yeast. The structure and related experiments show in detail an essential point of Cse4 contact and provide information about the arrangement of the inner kinetochore.

## Introduction

Kinetochores assemble at centromeres by interacting with a variant histone H3 known as centromere protein A (CENP-A) in vertebrates and as Cse4 in *Saccharomyces cerevisiae* and other point-centromere yeast. CENP-A and Cse4 deviate from canonical histone H3 at residues interspersed in the histone core and in the sequence and length of the N-terminal segment (Fig. 1A). Components of the assembled kinetochore that recognize these CENP-A-or Cse4-specific features include yeast Mif2 (CENP-C in vertebrates) and one or more subunits of the yeast Ctf19 complex (Ctf19c), which is homologous to the human CCAN (for Constitutive Centromere Associated Network; Fig. 1B). An important open question is how these factors “read” distinguishing features of Cse4 to restrict kinetochore assembly to centromeres.

**Figure 1.**
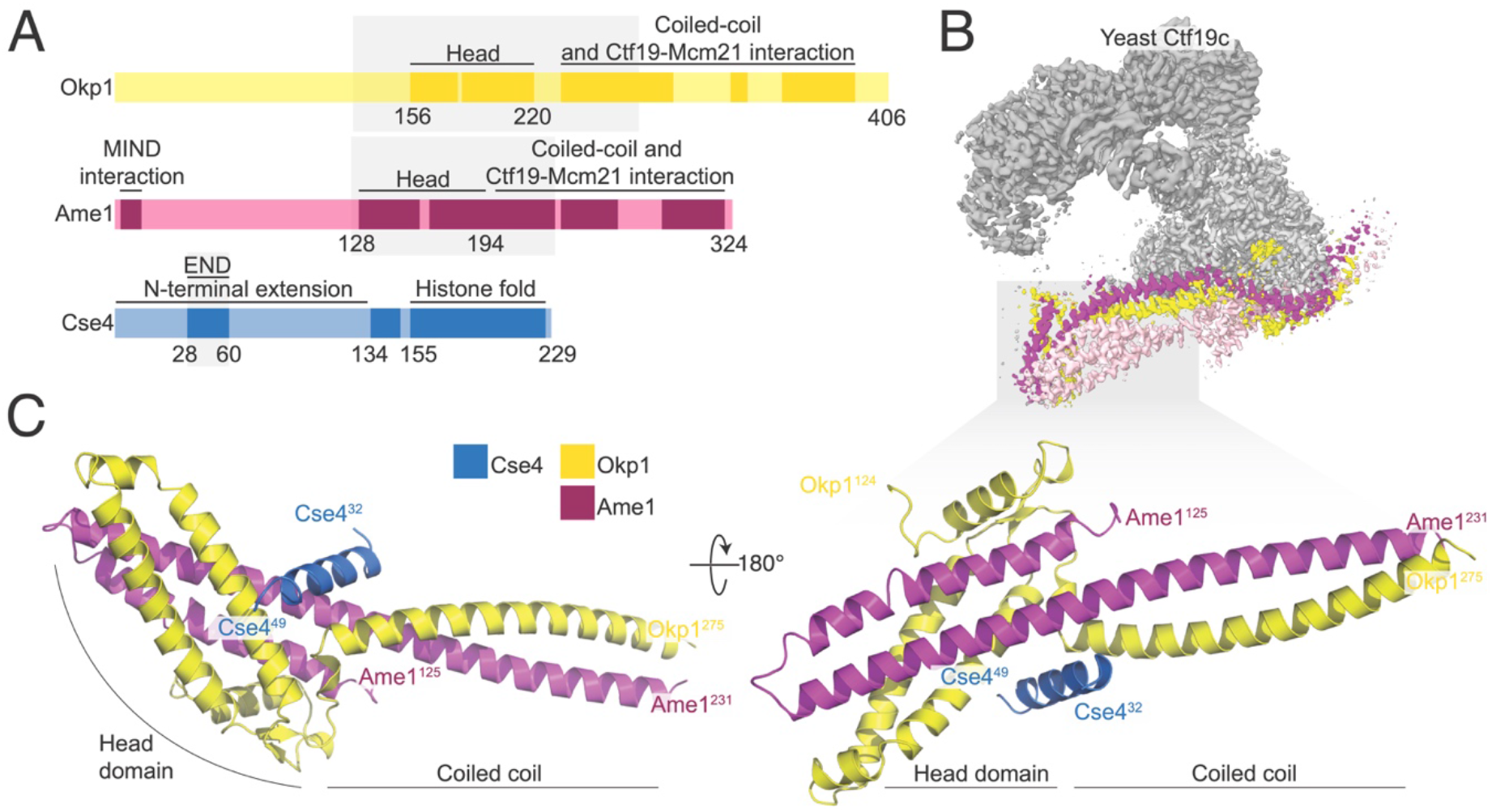
(A) Domain diagram showing the relevant regions of Okp1, Ame1, and Cse4. The shaded boxes demarcate the minimal Okp1-Ame1 peptides used for crystallography. The “Head” domains correspond to the previously described four-helix bundle (by analogy to the MIND complex; (*30*)). An N-terminal peptide of Ame1 connects to spindle microtubules indirectly via the MIND complex (*8*). (B) Composite cryo-EM structure of the yeast Ctf19c with the Okp1-Ame1 complex colored yellow (Okp1) and magenta (Ame1). Nkp1 and Nkp2 are colored light pink. (C) Crystal structure of the Okp1-Ame1-Cse4^END^ complex. Protein chains are colored as in panel B. Cse4 is blue.

The Cse4 N-terminal extension is much longer than its histone H3 counterpart (∼140 and ∼40 amino acid residues, respectively). Despite its length, a relatively short segment of the Cse4 N-terminal extension is necessary and sufficient for cell growth (residues 28 to 60, Fig. 1A) (*1*). Accordingly, this segment was named the Cse4 <u>E</u>ssential <u>N</u>-terminal <u>D</u>omain (Cse4^END^). Early yeast two-hybrid experiments suggested that Ctf19 binds Cse4^END^ (*1*). More recent biochemical reconstitution experiments showed that the Okp1-Ame1 heterodimer, which is a Ctf19c sub-module that binds the Ctf19 protein, is the direct binding partner of Cse4^END^ (*2-4*).

Okp1 and Ame1 are the only fully essential components of the Ctf19c. They associate as an elongated heterodimer with a globular “head” and an extended alpha-helical coiled-coil “shaft.” Loops in both chains interrupt the shaft and interact with Ctf19 and Mcm21 to make the COMA complex (*4-6*). Nkp1 and Nkp2 bind along the length of the Okp1-Ame1 dimer, making a six-protein complex (*7*). Okp1, Ame1, Ctf19, and Mcm21 all have long, flexible N-terminal extensions. Only that of Ame1 is essential; it indirectly recruits the microtubule-binding Ndc80 complex to the kinetochore, thus enabling chromosomal contact with the mitotic spindle (*8*).

Molecular recognition of Cse4 by the Okp1-Ame1 heterodimer has not been resolved, despite recent cryo-EM structures that show in detail nearly all protein contacts that contribute to inner kinetochore assembly. An experimental solution will constrain models for centromeric nucleosome binding by the Ctf19c. There are also several reported post-translational modifications of the Cse4 N-terminal extension that are thought to control kinetochore function (*2, 9-15*). Cse4 binding to Okp1-Ame1 can be reconstituted *in vitro* (*2-4*), but a published cryo-EM structure that includes Cse4 and the assembled Ctf19c did not show Cse4^END^ density (*16*). In our own efforts to solve this problem, we have attempted to determine the structure of the Ctf19c bound either to a reconstituted Cse4 nucleosome particle or to a minimal Cse4^END^ peptide, but the resulting maps do not show density corresponding to Cse4^END^. In the current work, we have used X-ray crystallography to identify the Cse4^END^ binding site.

We show here that Cse4^END^ binds at the head-shaft junction of Okp1-Ame1. We determined a 1.8 Å resolution crystal structure of a truncated Okp1-Ame1 bound with a Cse4^END^ peptide to specify in detail the amino acid residues involved, and we have shown by binding experiments and yeast genetics that the interaction is functionally relevant in cells.

## Results

### The crystal structure of Cse4^END^ with Okp1-Ame1

We determined the crystal structure of the Cse4^END^ peptide bound to a minimal Okp1-Ame1 heterodimer (Fig. 1A). Crystals containing all three proteins, including the full Cse4^END^ sequence, grew in space group 4_2_2_1_2 and yielded diffraction data to a minimum Bragg spacing of 1.8 Å. We determined the structure by molecular replacement (MR) as described in the Methods section and in Table S1.

In the crystal structure of the heterotrimeric complex, Cse4^END^ binds at the junction between the globular head of Okp1-Ame1 and the proximal part of the coiled-coil shaft. Most ordered Cse4 residues in the crystal structure have α-helical character (residues 34-46), and there are short flexible extensions (residues 32-34 and 46-49). The Cse4^END^ position is essentially as predicted by AlphaFold 2 ((*17, 18*) and this work; Fig. S1A-B),). Nkp1 and Nkp2 bind an adjacent site in the assembled Ctf19c, but it is not clear from comparison of the cryo-EM and crystal structures (Fig. S1C) whether they would need to partly dissociate from Okp1-Ame1 to accommodate Cse4^END^. Aside from Nkp1 and Nkp2, the Cse4^END^ binding site is exposed and on an external surface in structures of the assembled Ctf19c (*4, 16*).

The orientation of the four-helix bundle that makes up the Okp1-Ame1 head domain differs from what we and others observed in cryo-EM reconstructions of the Ctf19c (Fig. S1C) (*4, 16*). In the current structure, the Okp1-Ame1 head tilts away from the shaft. This is most evident for Ame1, where there is no discernable break in the extended helix that connects the shaft and head, whereas the same helix bends at Ame1-I195 in the previous Ctf19c structures.

### Specific contacts that enable Cse4^END^-Okp1-Ame1 binding

The three-way interface between Cse4, Okp1, and Ame1 has a hydrophobic core surrounded by strong polar interactions. The Cse4^END^ residues at the interface are conserved across point-centromere yeast, as are most of their partners in Okp1 and Ame1 (Fig. 2A, S2). Hydrophobic contacts between Cse4 and Okp1 include Cse4-L42 and Okp1-I234 (Fig. 2B). Likewise, Cse4-L41 and -I34 contact Ame1 I195 (Fig. 2C). Peripheral charged contacts include an electrostatic interaction between Cse4-R46 and Okp1-E235. Cse4-R37 makes complementary charge contacts with Ame1-D191 and -D194. Both contacts are conserved in yeast. This overall arrangement, with multiple charged contacts surrounding a hydrophobic core encompassing all three proteins, explains how a short Cse4^END^ peptide binds the Okp1-Ame1 dimer with an equilibrium dissociation constant of ∼90 nM (see below).

**Figure 2.**
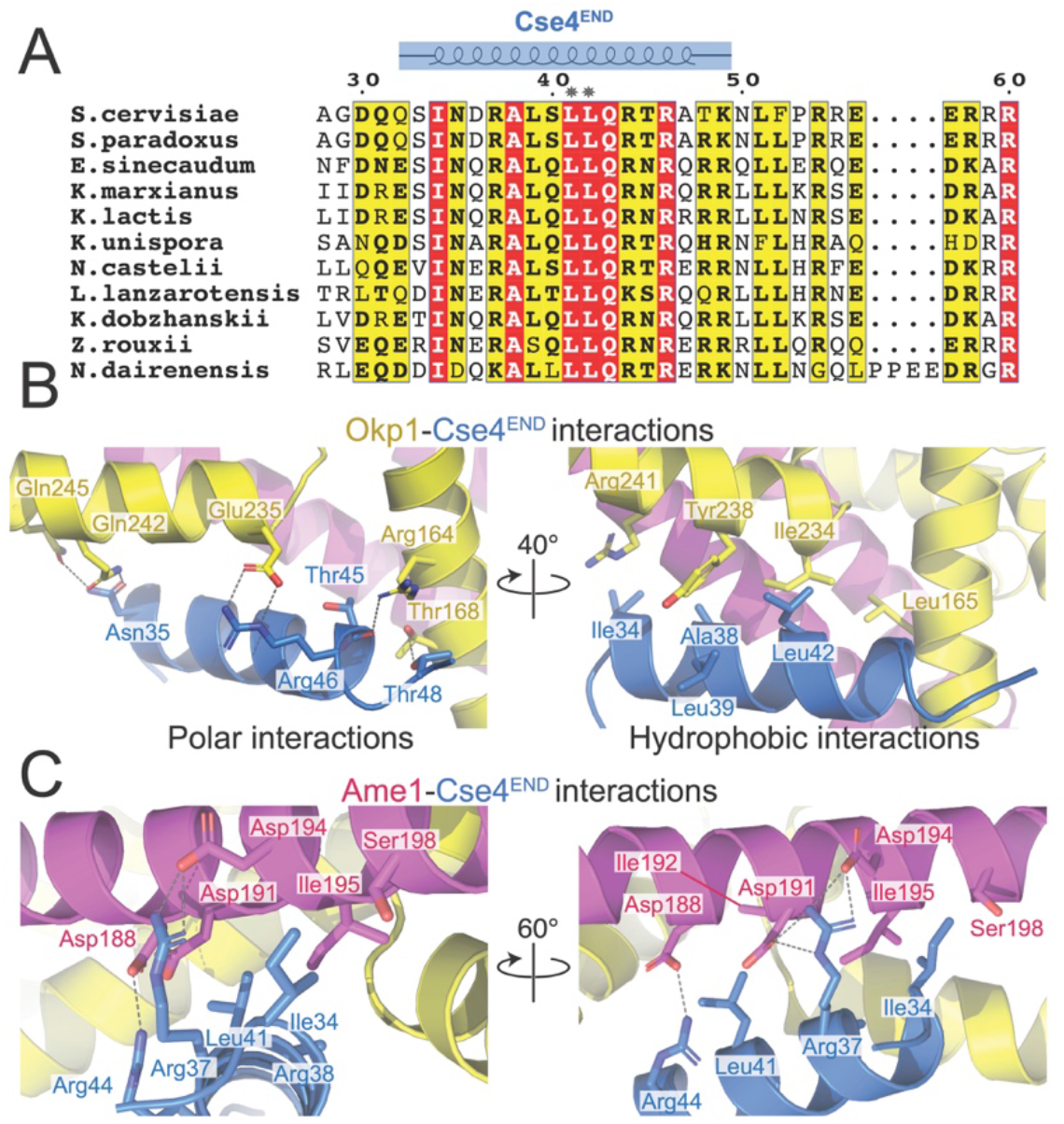
(A) Protein sequence alignment showing the Cse4^END^ peptide. Numbering corresponds to the *S. cerevisiae* Cse4 protein. Amino acid residues 28 to 60 are shown. The blue diagram above shows the residues visible in the crystal structure. Asterisks mark L41 and L42. (B) Close-up views of the Okp1-Cse4 interface. Polar interactions are shown on the left. Hydrophobic contacts are shown on the right. (C) Two views of the Ame1-Cse4 interface.

Comparison with Cryo-EM structures of the Ctf19c shows that Nkp1 and Nkp2 bind close to the Cse4^END^ binding site on Okp1-Ame1. To test whether Cse4^END^ binding depends on Nkp1-Nkp2, we used pulldown assays to reconstitute the interaction between Cse4^END^ and Okp1-Ame1. Consistent with published data (*2-4*), a glutathione-S-transferase (GST) fusion of Cse4^END^ bound Okp1-Ame1 (Fig. 3A). GST-Cse4^END^ bound the four-protein COMA complex and the six-protein COMA-Nkp1-Nkp2 complex equally well. Truncation of Okp1-Ame1 to the minimal heterodimeric complex used for crystallization (Okp1^125-275-^Ame1^124-231^) preserved GST-Cse4^END^ binding (Fig. S3A). To supplement the pulldown assays and to quantify Cse4^END^ binding to Okp1-Ame1, we created a fluorescence polarization (FP) assay using a fluorescent Cse4^END^ peptide (Fig. 3B) An equilibrium dissociation constant of ∼90 nM describes the binding event, and this value matches one previously reported (*2*). Adding excess Nkp1-Nkp2 protein to the binding reaction did not change the dissociation constant, consistent with the pulldown results. Therefore, Cse4 can bind the assembled COMA-Nkp1-Nkp2 complex.

**Figure 3.**
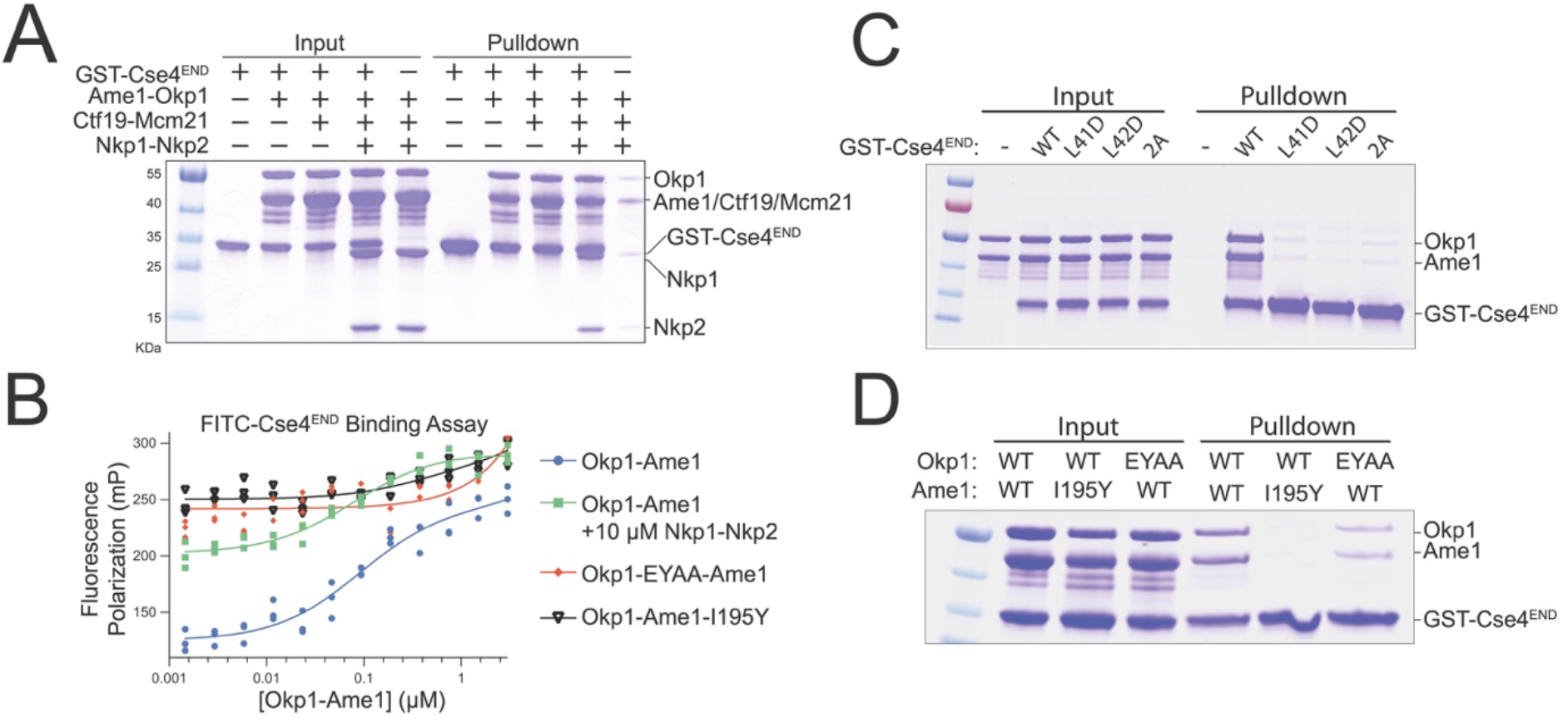
(A) GST pulldown assay for Cse4 binding. The indicated COMA complex proteins were tested for their association with immobilized GST-Cse4^END^. (B) Fluorescence polarization assay for quantification of Cse4 binding affinities. FITC-labeled Cse4^END^ peptide was incubated with increasing concentrations of the indicated protein complexes. See also Table 1. (C-D) GST pulldown assays as in panel (A). Cse4^END^ and its mutants (L41D, L42D, or L41,42A) were tested in panel (C). The indicated Okp1-Ame1 complexes were tested for their association with GST-Cse4^END^ in panel (D).

### Effects of mutations on affinity of Cse4^END^ for Okp1-Ame1

We tested the importance of residues at the interface between Cse4^END^ and Okp1-Ame1 using the biochemical assays described above. The *cse4-L41D, cse4-L42D*, and *cse4-L41,42A* (*cse4-2A*) mutations disrupt central hydrophobic interactions with Okp1-Ame1. The corresponding Cse4^END^ peptides failed to bind recombinant Okp1-Ame1 in the pulldown assay (Fig. 3C). The same was true in a pulldown assay carried out with the minimal Okp1-Ame1 complex used for crystallography (Fig. S3B). These experiments confirm the importance of hydrophobic contacts at the core of the three-protein interface.

**Table 1.** Affinity measurements for Cse4^END^ binding determined by fluorescence polarization. Full length Okp1-Ame1 complex was used. Equilibrium constant values are given in nM ± 95% confidence interval (*K*_D_ – equilibrium dissociation constant, r^2^ – goodness of fit, n.d. – not determined).

| | Okp1-Ame1 version | $K_D$ | $r^2$ |
| --- | --- | --- | --- |
| Okp1-Ame1 (WT) | FL | 91 ± 25 nM | 0.9481 |
| Okp1-Ame1 W(T)<br>+ 10 μM Nkp1-Nkp2 | FL | 80 ± 17 nM | 0.9401 |
| Okp1(EYAA)-Ame1 | FL | > 3 μM | n.d. |
| Okp1-Ame1(I195Y) | FL | >> 3 μM | n.d. |

We used the crystal structure to create Okp1-Ame1 mutations that specifically interfere with Cse4^END^ binding. Among those tested, *ame1-I195Y* and *okp1-E235,Y238A* (*okp1-EYAA*) were the most potent disruptors of Cse4 binding. Both mutations prevented Cse4^END^ binding in the pulldown and fluorescence polarization assays (Fig. 3B,D). They also prevented Cse4^END^ binding when introduced into the minimal Okp1-Ame1 complex (Fig. S3C). In addition to these mutations, we also tested the *ame1-D191A* and *-D194A* mutations, which prevent coordination of the conserved Cse4-R37 via hydrogen bonding interactions. Neither mutation perturbed Cse4^END^ binding, and the combination (*ame1-DDAA*) disrupted Cse4 binding in the pulldown assay, but only slightly (Fig. S3D).

### Effects of Cse4^END^ binding mutants on cell growth

To test whether Cse4^END^ mutations that disrupt Okp1-Ame1 binding interfere with cell division, we used a plasmid shuffling assay in which a plasmid-borne complementing *CSE4* allele restores the viability of *cse4Δ* cells. Selection against the complementing allele reveals the phenotype associated with a *CSE4* test allele carried on a second plasmid. The *cse4-2A* test allele did not support cell growth, and the *cse4-L41A* (*cse4-1A*) test allele produced slower-growing cells than did the *CSE4* allele (Fig. 5A). Therefore, Cse4^END^ hydrophobic residues required for Ame1-Okp1 binding are also required for cell viability.

We next tested whether Okp1-Ame1 residues that interact with Cse4^END^ are essential. To do so, we sporulated diploid cells heterozygous for the *ame1-I195Y* and *okp1-E235A,Y238A* mutations and examined their haploid progeny (Fig. 4B-D). Neither mutation produced obvious dominant defects in the parental heterozygous cells, and the expression levels of these mutant proteins matched their wild type counterparts (Fig. S4A). Both mutations permitted cell growth in haploid spores but were lethal in combination with *mcm21Δ* (Fig. 4C). This result matches the previous observation that non-lethal Cse4^END^ mutations are synthetic-lethal with Ctf19c mutations (*e*.*g*., *cse4-R37A chl4Δ* double mutant cells are inviable) (*2, 15*). If *ame1-I195Y* and *okp1-E235A,Y238A* each support weak Cse4 binding below the detection limit of our biochemical assay, then introduction of both mutations might completely ablate the interaction, yielding inviable spores. To test this idea, we examined *okp1-E235A,Y238A ame1-I195Y* double mutants and found that these spores produced tiny colonies (Fig. 4D). We carried out the same experiments for the *ame1-D191A,D194A,I195Y* and *okp1-I165A,I234A* mutations and observed the same set of double mutant phenotypes in progeny spores (Fig. S4B-F).

**Figure 4.**
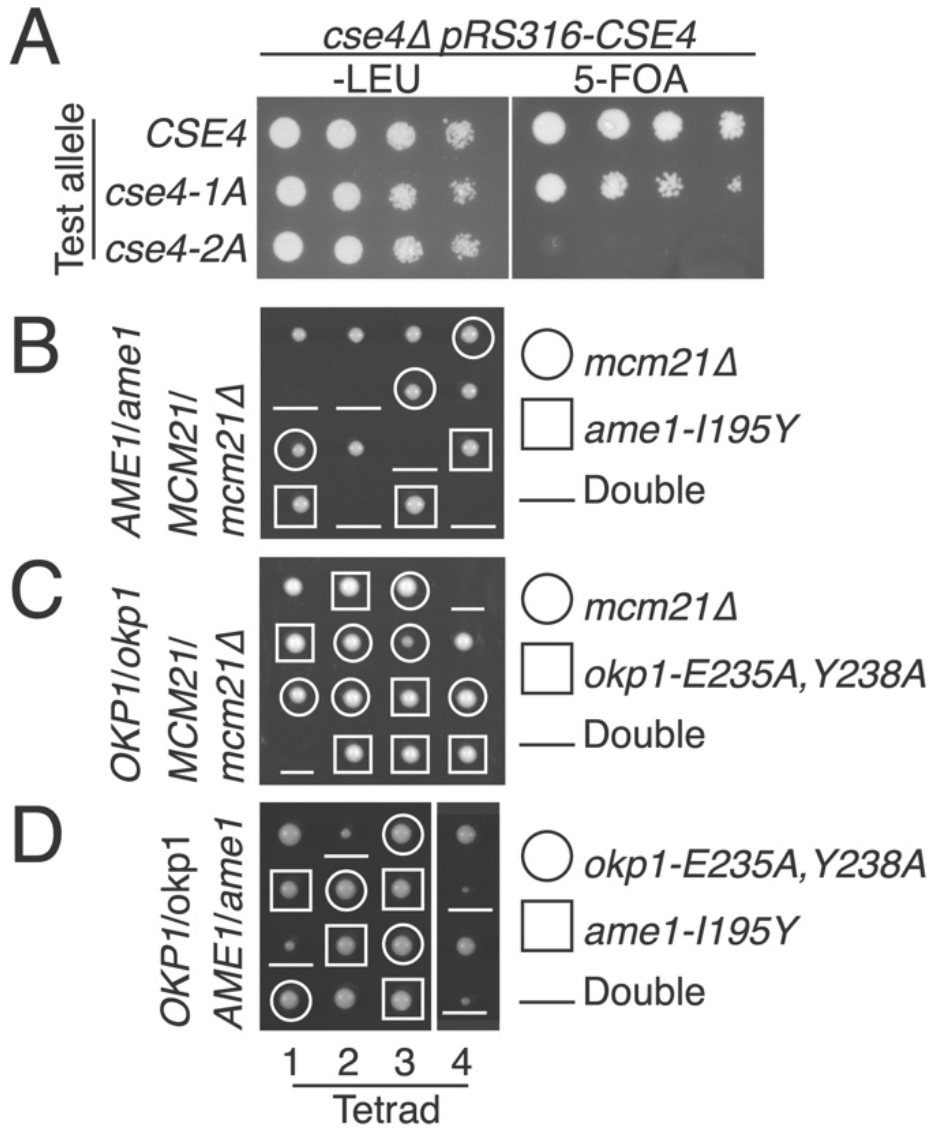
(A) Growth of *cse4* mutants. Loss of a complementing *CSE4* plasmid (*pRS316-CSE4*) was induced by growth on 5-FOA. Test *cse4* alleles are indicated at left (*cse4-1A* – L41A; *cse4-2A* – L41,42A). (B-D) Haploid progeny from sporulation of the indicated heterozygous diploid strain (left). Spore genotypes and mutant alleles are given at right. Meiotic products from each tetrad are arranged vertically.

Therefore, the Cse4 binding site on Okp1-Ame1 is required for cell viability. As demonstrated by the synthetic lethality when combined with *mcm21Δ*, an intact Ctf19c can compensate for partial disruption of the Cse4 binding site *in vivo*.

## Discussion

We have determined the crystal structure of an essential segment of Cse4 bound to the inner kinetochore proteins Okp1 and Ame1. This segment, Cse4^END^, binds the junction between the Okp1-Ame1 coiled-coil shaft and head domains. Minimal Okp1-Ame1 mutations that disrupt Cse4 binding (this work) or MIND binding (*8*), are lethal, indicating that the main essential function of Okp1-Ame1 is to connect centromeric DNA (via Cse4) to spindle microtubules (via MIND-Ndc80). Force transmission through these connections has been reconstituted in a minimal biochemical system (*19*). Whether this axis is the essential conduit for the force that moves chromosomes *in vivo* or if its essential function is regulatory remains to be seen.

The Cse4-Okp1-Ame1 structure constrains models for centromeric nucleosome recognition. An initial Ctf19c structure in the absence of the Cse4 nucleosome suggested a straightforward docking model for Ctf19c-Cse4 binding, with which the observed position of Cse4^END^ in the current structure is compatible (*4*). This model requires repositioning of the Ctf3c-Cnn1-Wip1 heptamer and the Okp1-Ame1 head domains, along with partial DNA unwrapping from the nucleosome itself. Flexibility of the Ctf3 module and partial DNA unwrapping have been observed (*17, 20, 21*), and we report that the Okp1-Ame1 head domain is indeed mobile with respect to the adjacent coiled-coil shaft.

More recent Cryo-EM structures of the yeast inner kinetochore (*16, 17*) have raised the possibility of alternative Ctf19c-nucleosome poses. These rely principally on contact between Ctf19c proteins and the DNA phosphate backbone. In one case, the Cbf1 protein contributes to DNA sequence specificity (*17*). In these published structures, a ∼60 Å peptide linker would be required to connect the Cse4 histone core to the Cse4^END^ peptide. This contradicts genetic evidence, which shows that fusion of the Cse4^END^ segment directly to the histone core (at residue 130) supports cell viability at a level indistinguishable from that of a wild type *CSE4* allele (*1*). An alternative assembly, in which an extra and presumably essential COMA complex orbits the Ctf19c and attaches through Cse4^END^ would satisfy this discrepancy (*17*), but there is at present no evidence for such an arrangement. Instead, we suggest that either *i*) the heterotrimeric contacts we have described serve a cell cycle-specific function and do not occur in the fully assembled Ctf19c or *ii*) the published Ctf19c-Cse4 structures do not recapitulate the physiologically relevant positions of the centromeric histone proteins and of the centromere itself.

This work serves to re-emphasize the question, what distinguishes the structure of the assembled Ctf19c, which is known, from its structure when bound to the centromeric nucleosome? Neither the published Ctf19c-Cse4 structures nor our own unpublished structures, in which we have attempted to visualize just the Cse4^END^ peptide bound to the assembled Ctf19c, show any discernable Cse4^END^ density. That Cse4^END^ binds recombinant COMA but not the assembled Ctf19c suggests a structural switch accompanies nucleosome engagement. The altered orientation of the Okp1-Ame1 head domain in the current structure is one candidate for such a switch. Relevant post-translational modifications of Ctf19c proteins, Cse4, and Mif2 have all been reported (*2, 9, 22, 23*). It will be important to determine which modifications are required for inner kinetochore assembly and under what cell cycle conditions they contribute.

## Acknowledgements

We thank Cindy Zhang for technical assistance. X-ray diffraction data were recorded at beamline ID-24-E (operated by the Northeast Collaborative Access team: NE-CAT) at the Advanced Photon Source (APS, Argonne National Laboratory). We thank the staff of the NE-CAT beamlines for help in data collection. NE-CAT is supported by NIH grant P30 GM124165, using resources of the APS, operated by Argonne National Laboratory under Contract DE-AC02-06CH11357. JC and HZ are supported by RO1 GM116897. SCH is an Investigator in the Howard Hughes Medical Institute.

## Methods

### Protein Expression and Purification

Heterodimers of full length Okp1-Ame1, Ctf19-Mcm21, and Nkp1-Nkp2 were expressed and purified in *E*.*coli* as previously described (*4*). Briefly, transformed cells were grown to an OD_600_ of 0.5 at 37 °C in 2XYT media with antibiotic selection, induced with 0.5 mM of Isopropyl β-D-1-thiogalactopyranoside (IPTG) at 18°C for ∼16 h, and harvested by centrifugation for 20 minutes at 4000 rpm. Pelleted cells were lysed by sonication in the lysis buffer containing 25 mM HEPES, pH 7.5, 800 mM NaCl, 10 mM imidazole,1 mM tris(2-carboxyethyl)phosphine (TCEP), 0.1 mg/mL PMSF, 5 mM β-mercaptoethanol, 1 mM PMSF, 1 μg/mL pepstatin, 1 μg/mL aprotinin, 1 μg/mL leupeptin, 30 μg/mL DNase I, and an EDTA-free protease inhibitor tablet (Roche, cOmplete™). After sonication, lysate clarified by centrifugation was passed over TALON Metal Affinity Resin (Takara), washed with 10 column volumes (CV) of lysis buffer, and eluted with 400 mM imidazole. The eluate was further purified on a HiTrap SP or Q ion-exchange column (Cytiva). The peak fractions were pooled for gel filtration in 20 mM Tris, pH 8.5, 200 mM NaCl, 1 mM TCEP. Peak fractions were collected, concentrated, snap frozen, and stored at −80 °C until use.

For pulldown assays, Cse4 residues 28-60 (Cse4^END^) was cloned into a pLIC vectors with a TEV-cleavable N-terminal Glutathione S-transferase tag and transformed into Rosetta™ 2 (pLysS) competent cells (Sigma-Aldrich). GST-Cse4^END^ was purified as described above, using Pierce Glutathione Agarose (Thermo Scientific) and eluting with L-Glutathione.

To find a minimal dimeric construct of Okp1-Ame1 that could bind Cse4, we cloned various truncated constructs of Okp1 and Ame1 genes into the pet-Duet bacterial expression vector, with a poly-histidine tag and tobacco etch virus (TEV) protease site at the N terminus of Okp1 and no tag on Ame1. We found that Okp1^125-275^ -Ame1^124-231^ can be expressed in Rosetta™ 2 (pLysS) competent cells (Sigma-Aldrich) and purified as described above, except that the N-His_6_ tag on Okp1 was cleaved with TEV protease at 30°C for one hour before the ion-exchange step. Final proteins after gel filtration were concentrated to ∼ 15 mg/ml, snap frozen, and stored in 100 mM HEPES, pH 7.5, 300 mM ammonium sulfate, and 1 mM TCEP, at -80°C until use.

### Pulldown assays

GST-Cse4^28-60^ was mixed with Okp1-Ame1, COMA, COMA-Nkp1-Nkp2, or Okp1^125-275^ - Ame1^124-231^ and the mixed samples were incubated with glutathione agarose resin at 4°C for 1 hour in binding buffer containing 20 mM Tris, pH 8.5, 200 mM NaCl and 1 mM TCEP. Resin was washed at least three times with the same buffer to remove unbound protein, and bound protein then eluted with glutathione. Samples of input and eluate were collected for analysis by SDS-PAGE.

### Fluorescence Polarization (FP) Assays

FITC-Cse4^28-60^ peptide was synthesized by the Tufts University Core Facility. 20 nM of FITC-Cse4^28-60^ peptide was used in all reactions, and Okp1-Ame1 concentrations were varied to determine the dissociation constant (*K*_*D*_), in the reaction buffer containing 100 mM HEPES, pH 7.5, 200 mM NaCl, 1 mM TCEP. The reactions were analyzed after incubating for 30 min at room temperature. FP readings were recorded with a Perkin Elmer EnVision (ICCB-Longwood Screening Facility, Harvard University) and each curve was repeated in triplicate. GraphPad Prism was used for all data fitting.

### Protein Crystallization and diffraction data collection

Purified Okp1^125-275^ - Ame1^124-231^ was mixed with peptide Cse4^28-60^ (synthesized at the Tufts University Core facility) in a ratio of 1: 1.3 and diluted to 9 mg/ml. The best crystals were grown within ∼3 days by hanging drop vapor diffusion at 18 °C against reservoir solution containing 19% PEG 8000, 0.55 M lithium sulfate and a 1:1 sample:well drop ratio. Crystals were transferred into mother liquor supplemented with 25% glycerol and flash-frozen in liquid nitrogen. Diffraction data to 1.8 Å minimum Bragg spacing were collected on beamline 24-ID-E at the Advanced Photon Source (Argonne National Laboratory). The complex crystallized in space group P 42 21 2 (a = 154.52, b = 154.52, c = 37.18). Data collection statistics are in Table S1.

### Structure determination

X-ray diffraction data processing was carried out with xia2 (https://xia2.github.io/index.html) using DIALS (*24*)and Aimless (*25*). The Okp1^162-275^ - Ame1^124-231^ structure from the yeast the Ctf19c/CCAN structure (PDB: 6NUW) failed to yield a molecular replacement solution in Phenix(*26*), but the AlphaFold2 (*18*) prediction for Okp1^125-275^ - Ame1^124-231^ yielded a robust solution with clear density for the Cse4 peptide. Model building was carried out in Coot (*27*) and refined in Phenix(*26*). Coordinates and diffraction data have been deposited in the protein data bank (PDB ID: 8T0P). Refinement statistics are in Table S1.

### Construction of plasmids and yeast strains

Centromeric plasmids (pRS315 and pRS316) were linearized by NotI and repaired by PCR products of Cse4 (ChrXI: 345945-347290), Ame1 (ChrII: 646026-647638) and Okp1 (ChrVII: 853524-855435) flanked by 34 bp homology via homologous recombination in yeast using standard yeast transformation method. Plasmids were rescued from successful transformants via electroporation, sequenced, and then used to make other plasmid derivatives. These plasmid derivatives, including insertion of 3xFlag, TAF tag (3xFlag-TEV-ProteinA), KanMX6 marker and later point mutants, were generated by combining PCR fragments with overlapping homology regions (> 33 bp), via homologous recombination in yeast. Specifically, a 3xFlag tag was inserted at an internal XbaI site in Cse4 (*28*) and then a KanMX6 marker was inserted in the promoter region of Cse4 (ChrXI: 347222) to generate HZE2719. A TAF tag followed by KanMX marker was appended to the C-terminus of Ame1 (HZE2663) or Okp1 (HZE3198). These plasmids were then used to generate various point mutants as indicated.

A diploid of W303 strain (HZY1079) was used to generate a heterozygous deletion of Cse4, Ame1 or Okp1 using a natMX6 marker. After introducing pRS316-Cse4, pRS316-Ame1, or pRS316-Okp1, the transformed cells were sporulated to obtain the haploid *cse4, ame1* and *okp1* null mutants that were kept alive by their respective complementing plasmids. PCR products of *ame1* and *okp1* point mutants (with C-terminal TAF tag and G418 marker) were amplified from the mutant plasmids using a high-fidelity DNA polymerase (SuperFi, Invitrogen) and transformed into the respective haploid *ame1* and *okp1* null mutants. Correct integrations were identified by resistance to G418 and 5-Fluoroorotic acid (5-FOA), and nourseothricin-sensitivity, and confirmed by sequencing the genomic DNA of Ame1 and Okp1. Standard yeast genetic methods, including mating, sporulation, and dissection with a Singer dissection microscope, were used to construct various double mutants, as described by Lichten M. (*29*). Plates were incubated at 30 °C for 2 or 3 days before taking images using a BioRad ChemiDoc MP imaging system. The spores were identified based on the selection markers used to create each mutant. Yeast strains used are summarized in Table S2. Plasmids used are summarized in Table S3.

### Plasmid shuffling using URA3/5-FOA

Haploid strains *cse4Δ::natMX6 pRS316-Cse4* (HZY2970), were transformed with the indicated plasmids bearing *cse4* mutants. Successful transformants were grown up in Sc-Leu media to an OD_600_ ∼1, normalized based on cell density, and then spotted on the indicated plates following 5-fold serial dilutions. 5-fluoroorotic acid (Sc supplemented with 0.1% 5-FOA treatment removes the complementing *URA* plasmid, exposing the phenotypes of *cse4* mutants. Plates were incubated at 30°C for 2 or 3 days before taking images using a BioRad ChemiDoc MP imaging system.

## Figures

**Table S1.** Crystallographic and model statistics for the Okp1-Ame1-Cse4 crystal structure.

| **Supplementary Table 1. Statistics for Cse4<sup>END</sup> - Okp1<sup>xtal</sup> - Ame1<sup>xtal</sup> crystal structure** |  |
| --- | --- |
| **Crystal Parameters** |  |
| Space group | P 42 21 2 (No. 94) |
| Unit cell dimension | 154.305 (90) |
|  | 154.305 (90) |
|  | 37.114 (90) |
| **Data collection** |  |
| Wavelength (Å) | 0.97918 |
| Resolution (Å) | 109.11 -1.73 (1.76-1.73) |
| Unique reflections | 47590 (2301) |
| R<sub>merge</sub> | 0.125 (4.729) |
| R<sub>meas</sub> | 0.128 (4.818) |
| R<sub>pim</sub> | 0.025 (0.915) |
| CC<sub>1/2</sub> | 1.000 (0.355) |
| I/σ | 16.9 (1.0) |
| Completeness | 99.99% (97.75%) |
| Redundancy | 14.2% (14.1%) |
| **Refinement** |  |
| R<sub>work</sub>/R<sub>free</sub> | 0.2067/0.2364 |
| R<sub>msd</sub> |  |
| Bonds (Å) | 0.017 |
| Angles (°) | 1.554 |
| Average B factors (Å<sup>2</sup>) |  |
| Protein | 47.39 |
| **Ramachandran statistics (%)** |  |
| Favored | 97.83% |
| Allowed | 100% |
| Outliers | 0 |
| MolProbity clash score | 5.0 |

**Table S2.** Yeast strains used in this work.

| Yeast strains |  |
| --- | --- |
| HZY988 | *mcm21Δ:Hyg*, MAT alpha, W303 |
| HZY1077 | MAT a, W303 (*ade2-1, can1-100, his3-1115, leu2-3, 112trp-1, ura3-1, RAD5+*) |
| HZY1078 | MAT alpha, W303 (*ade2-1, can1-100, his3-1115, leu2-3, 112trp-1, ura3-1, RAD5+*) |
| HZY1079 | Diploid by crossing HZY1078 with HZY1077 |
| HZY2869 | *ame1-2DA (D191A, D194A)-TAF:G418*, MAT a, W303 |
| HZY2870 | *ame1-I195Y-TAF:G418*, MAT a, W303 |
| HZY2871 | *ame1-triple (D191A, D194A, I195Y)-TAF:G418*, MAT a, W303 |
| HZY2872 | *okp1-E235A-TAF:G418*, MAT a, W303 |
| HZY2873 | *okp1-Y238A-TAF:G418*, MAT a, W303 |
| HZY2874 | *okp1-E235A, Y238A-TAF:G418*, MAT a, W303 |
| HZY2875 | *okp1-I165A, I234A-TAF:G418*, MAT a, W303 |
| HZY2876 | *okp1-I165A-TAF:G418*, MAT a, W303 |
| HZY2877 | Diploid by crossing HZY988 (*mcm21Δ:Hyg*) with HZY2869 (*ame1-2DA-TAF:G418*) |
| HZY2878 | Diploid by crossing HZY988 (*mcm21Δ:Hyg*) with HZY2873 (*okp1-Y238A-TAF:G418*) |
| HZY2879 | Diploid by crossing HZY988 (*mcm21Δ:Hyg*) with HZY2874 (*okp1-E235A, Y238A-TAF:G418*) |
| HZY2880 | Diploid by crossing HZY988 (*mcm21Δ:Hyg*) with HZY2875 (*okp1-I165A, I234A-TAF:G418*) |
| HZY2881 | Diploid by crossing HZY988 (*mcm21Δ:Hyg*) with HZY2876 (*okp1-I165A-TAF:G418*) |
| HZY2882 | Diploid by crossing HZY988 (*mcm21Δ:Hyg*) with HZY2872 (*okp1-E235A-TAF:G418*) |
| HZY2911 | Diploid by crossing HZY988 (*mcm21Δ:Hyg*) with HZY2870 (*ame1-I195Y-TAF:G418*) |
| HZY2912 | Diploid by crossing HZY988 (*mcm21Δ:Hyg*) with HZY2871 (*ame1-triple-TAF:G418*) |
| HZY2943 | *ame1-triple (D191A, D194A, I195Y)-TAF:G418*, MAT a, derived from HZY2912 |
| HZY2945 | *ame1-triple-TAF:G418*, MAT alpha, derived from HZY2912 |
| HZY2970 | *cse4Δ:NAT*, pRS316-G419-3xFlag-CSE4, MAT alpha, W303 |
| HZY2971 | *okp1-E235A, Y238A-TAF:G418*, MAT alpha, W303, derived from HZY2879 dissection |
| HZY2976 | *okp1-I165A, I234A-TAF:G418*, MAT alpha, W303, derived from HZY2880 dissection |
| HZY2977 | *okp1-E235A, Y238A-TAF:NAT*, MAT alpha, W303, derived from HZY2971 |
| HZY2978 | *okp1-I165A, I234A-TAF:NAT*, MAT alpha, W303, derived from HZY2876 |
| HZY2979 | Diploid by crossing HZY2943 (*ame1-triple-TAF:G418*) with HZY2977 (*okp1-E235A, Y238A-TAF:Nat*) |
| HZY2980 | Diploid by crossing HZY2943 (*ame1-triple-TAF:G418*) with HZY2978 (*okp1-I165A, I234A-TAF:Nat*) |
| HZY2981 | Diploid by crossing HZY2943 (*ame1-I195Y-TAF:G418*) with HZY2977 (*okp1-E235A, Y238A-TAF:Nat*) |
| HZY2982 | Diploid by crossing HZY2943 (*ame1-I195Y-TAF:G418*) with HZY2978 (*okp1-I165A, I234A-TAF:Nat*) |
| HZY2983 | *okp1-I165D-TAF:G418*, MAT a, W303 |
| HZY2984 | *okp1-I234D-TAF:G418*, MAT a, W303 |
| HZY2985 | *okp1-I165D, I234D-TAF:G418*, MAT a, W303 |
| HZY2986 | *okp1-E235A, Y238A-TAF:Nat*, MAT a, W303, derived from HZY2979 dissection |
| HZY2988 | *okp1-E235A, Y238A-TAF:Nat, ame1-triple-TAF:G418*, MAT alpha, W303, derived from HZY2979 dissection |
| HZY2989 | *okp1-I165A, I234A-TAF:Nat*, MAT a, W303, derived from HZY2980 dissection |
| HZY2990 | *okp1-I165A, I234A-TAF:Nat, ame1-triple-TAF:G418*, MAT a, W303, derived from HZY2980 dissection |
| HZY2999 | *ame1-I195Y-TAF:G418 okp1-E235A/Y238A-TAF:Nat*, MAT alpha, derived from HZY2981 |
| HZY3900 | *ame1Δ:NAT*, pRS316-*AME1*, MAT a, W303 |
| HZY3901 | *ame1Δ:NAT*, pRS316-*AME1*, MAT alpha, W303 |
| HZY3904 | *okp1Δ:NAT*, pRS316-*OKP1*, MAT a, W303 |
| HZY3905 | *okp1Δ:NAT*, pRS316-*OKP1*, MAT alpha, W303 |

**Table S3.**
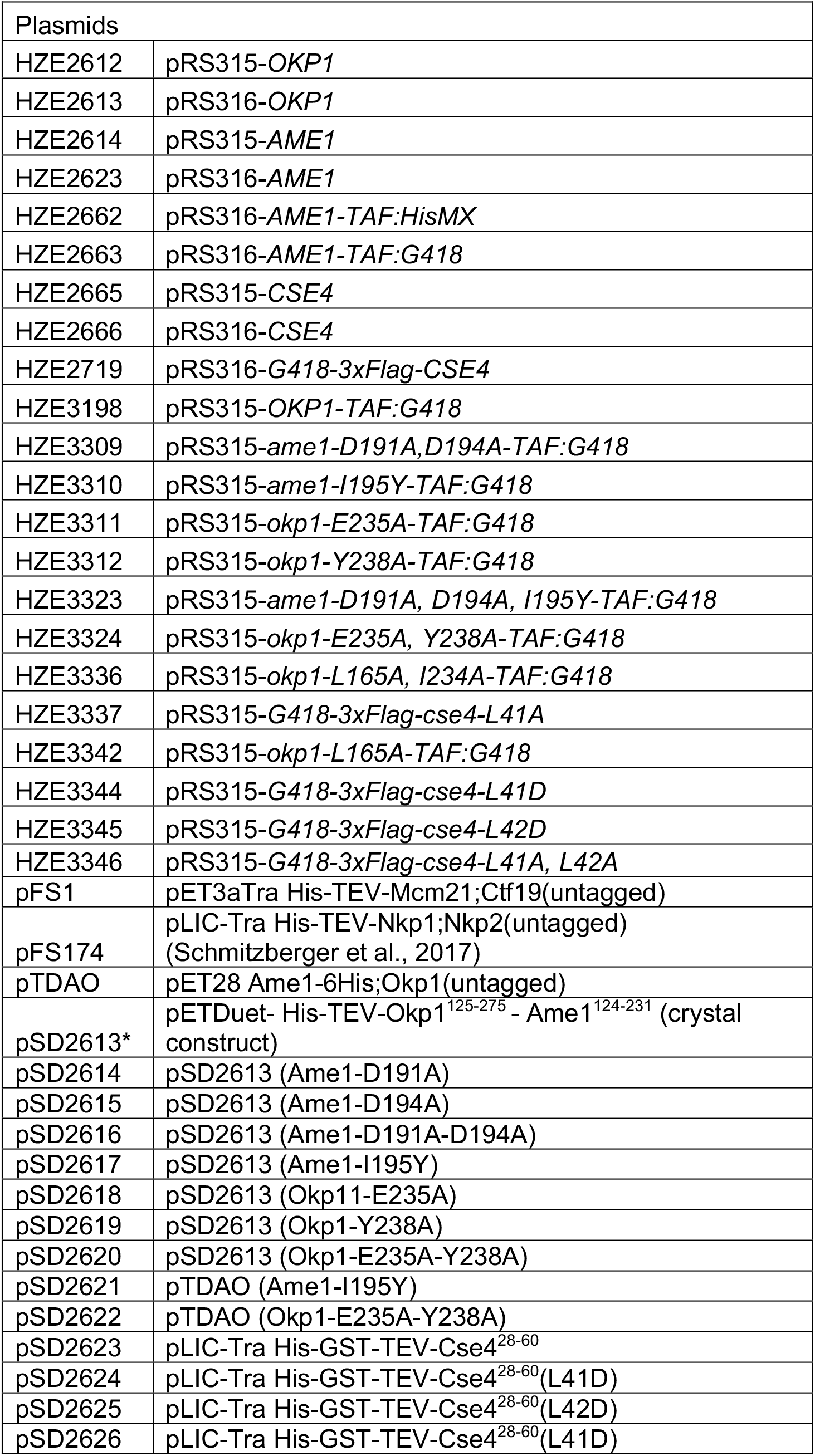
Plasmids used in this work.

**Figure S1.**
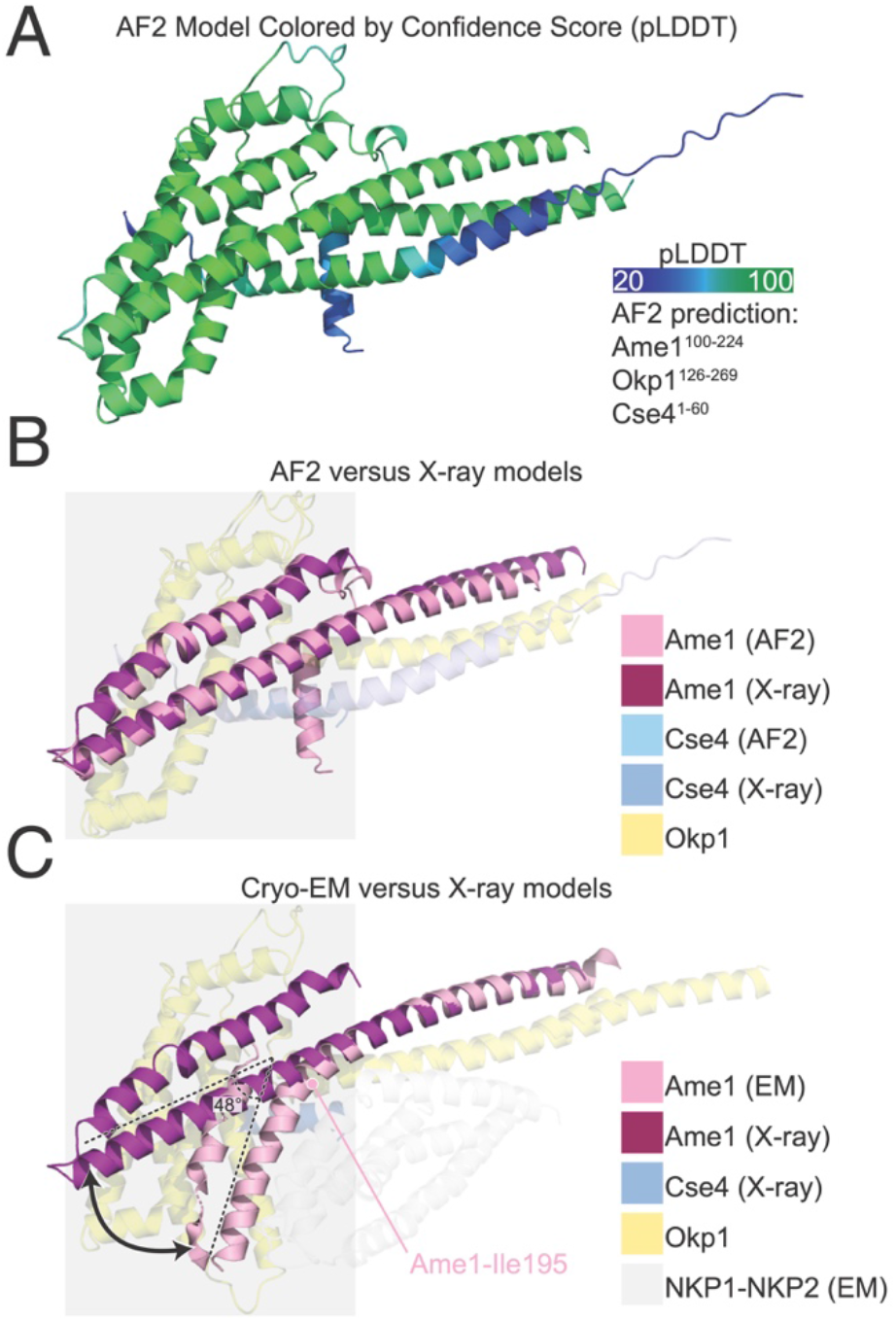
Flexibility at the Okp1-Ame1 head-coiled-coil joint and Nkp1-Nkp2 position. (A) The structure of Okp1-Ame1-Cse4 as predicted by AlphaFold 2 (AF2) (*18*). The model is colored according to confidence score (pLDDT) from low (blue) to high (green). The peptides used for prediction are given at right. (B) Overlay of Okp1-Ame1-Cse4 from AF2 with the current Okp1-Ame1-Cse4 crystal structure (X-ray). Only Ame1 (magenta and pink) is shown as an opaque chain for clarity. The gray box marks the Okp1-Ame1 head domain. (C) Overlay of the Okp1-Ame1-Cse4 structure from cryo-EM (EM) with the current crystal structure. The angle between the head and coiled coil shaft is indicated for the cryo-EM structure. Ame1-Leu195, which is the position at which Ame1 bends in the cryo-EM structure, is annotated. The Okp1-Ame1 head domain is marked as in panel B. Structures were aligned on the Okp1-Ame1 coiled coil shaft. The Nkp1-Nkp2 structure from cryo-EM (NKP1^2-76^; NKP2^4-84^) is shown as transparent gray chains.

**Figure S2.**
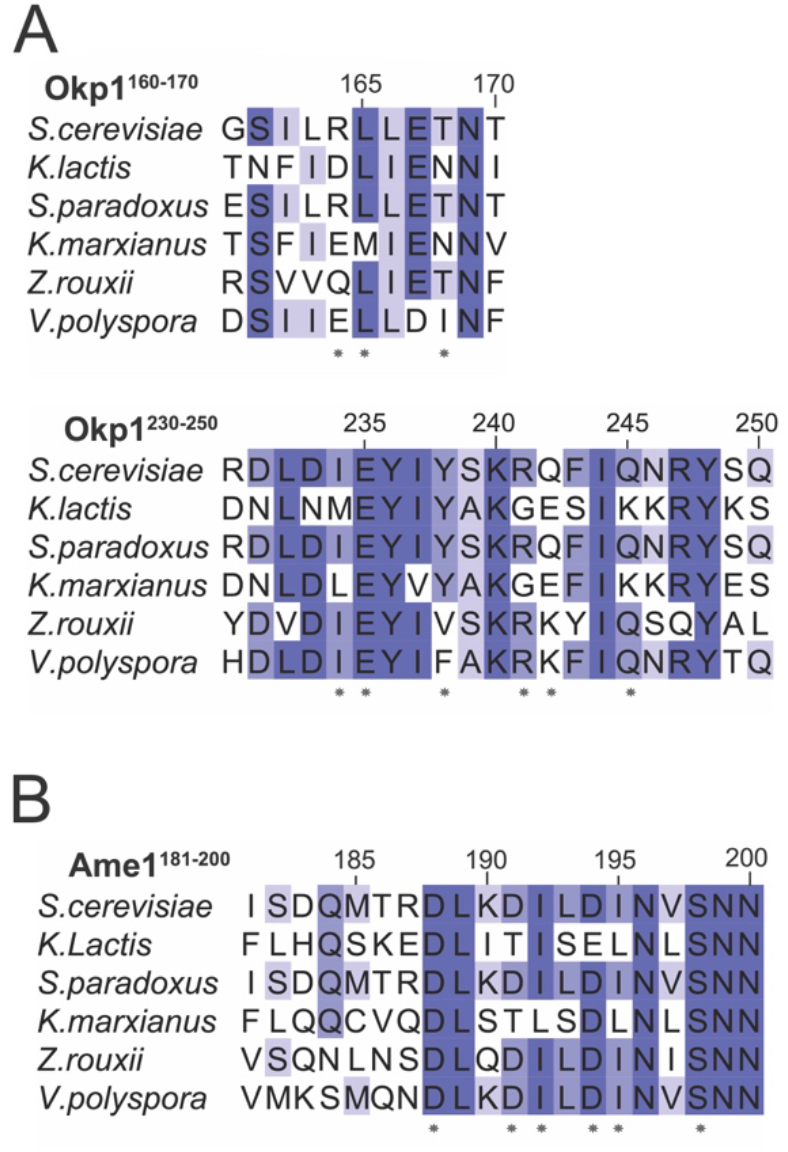
Protein sequence alignments for Okp1 and Ame1 covering the Cse4^END^ contacts shown in Figure 2.

**Figure S3.**
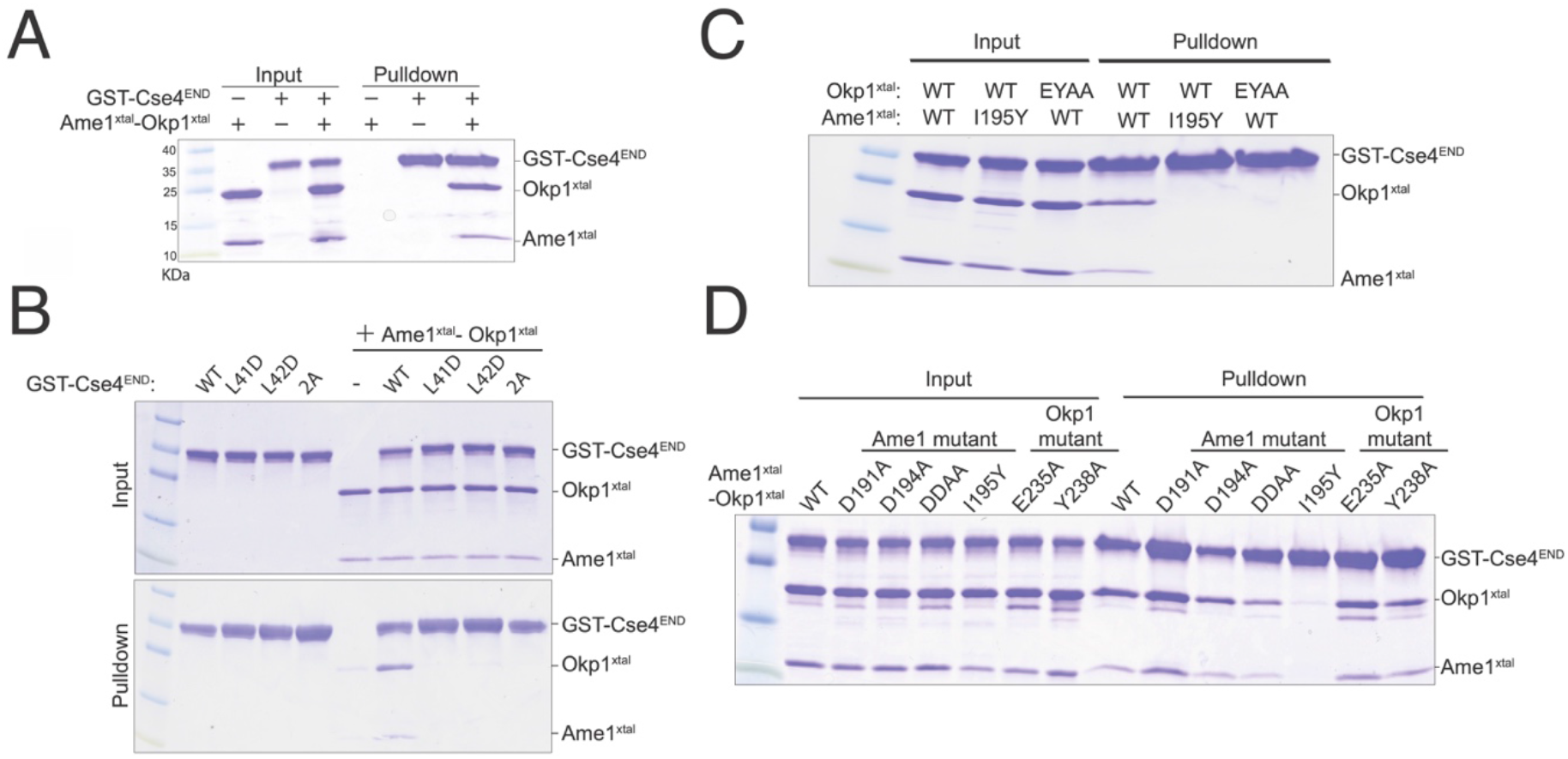
(A-D) The minimal Okp1-Ame1 complex used for crystallography was tested for its association with immobilized GST-Cse4^END^. (A) Pulldown assay showing binding between the minimal Okp1-Ame1 complex (Okp1^xtal^-Ame1^xtal^) and GST-Cse4^END^. (B) GST-Cse4^END^ and its mutants were tested for Okp1-Ame1 binding. (C) Various Okp1-Ame1 mutants (indicated above) were tested for binding to GST-Cse4^END^. (D) Various Okp1-Ame1 mutants were tested for Cse4^END^ binding as in panel (C).

**Figure S4.**
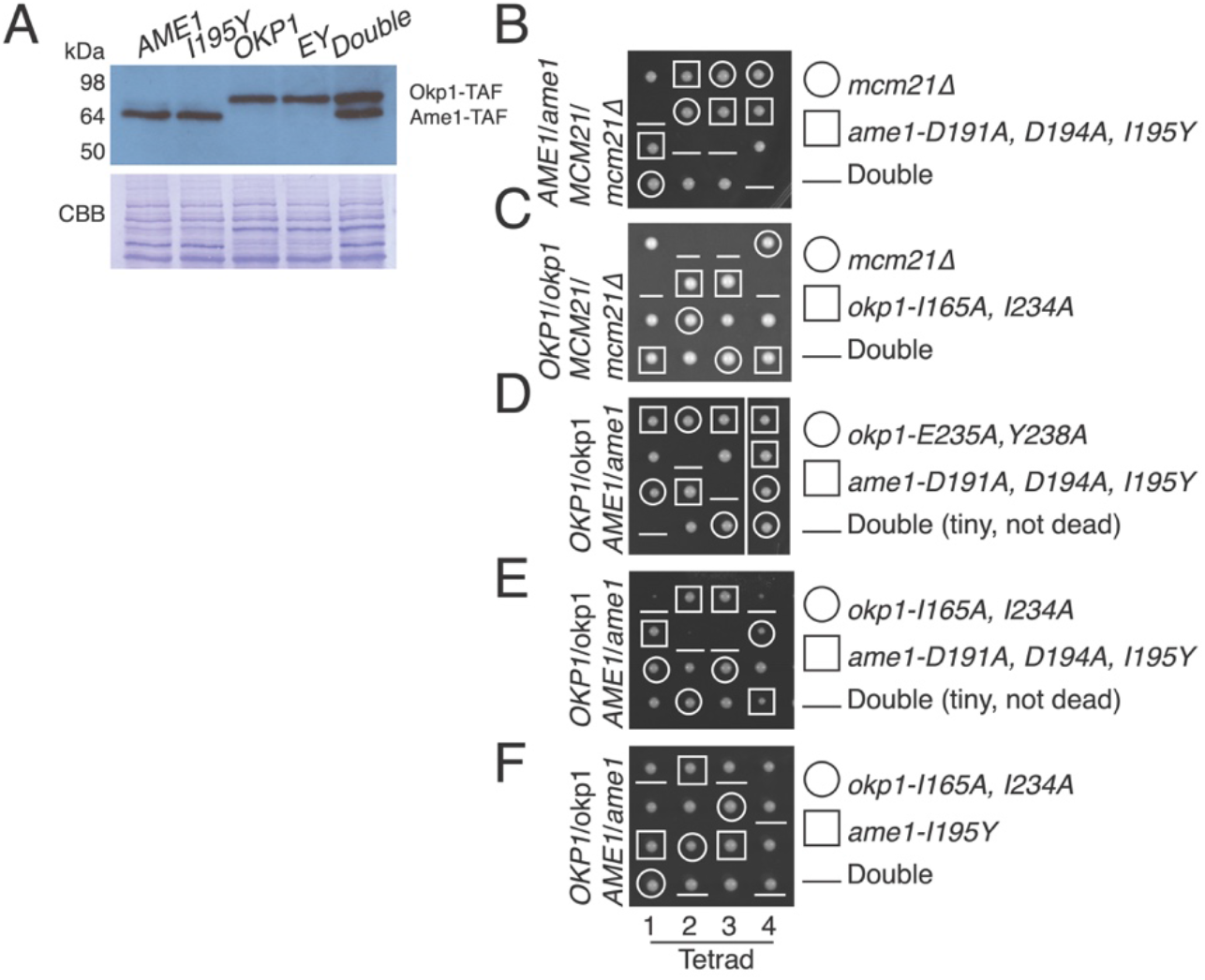
*In vivo* consequences of Okp1-Ame1 mutations. (A) Western blot showing expression of Ame1, Okp1, and their mutants in whole cell extracts (TAF – protein A-FLAG tag; anti-Protein A used for detection). (B-F) Tetrad dissection results as in Figure 4B-D. The mutants tested and the resulting spore genotypes are shown at right.

